# Transcriptomic analysis reveals coordinated metabolic reprogramming through activation of gluconeogenesis and suppression of fatty acid β-oxidation pathways during cold exposure in *Xenopus laevis*

**DOI:** 10.64898/2026.05.26.727745

**Authors:** Eiko Iwakoshi-Ukena, Makoto Suzuki, Megumi Furumitsu, Norisato Shimanoe, Yuki Narimatsu, Kazuyoshi Ukena, Hajime Ogino

## Abstract

**Background:** Ectothermic vertebrates exhibit substantial physiological plasticity in response to environmental temperature fluctuations. Among them, amphibians show particularly pronounced metabolic adjustments under cold conditions; however, the molecular mechanisms by which the liver adapts to low temperatures remain poorly understood. To address this gap, we investigated the hepatic transcriptional response to cold exposure in the African clawed frog (*Xenopus laevis*) using RNA sequencing (RNA-seq) and quantitative PCR.

**Results:** Exposure to 5°C for five days in *X. laevis* resulted in pronounced hyperglycemia and extensive transcriptional reprogramming in the liver. RNA-seq analysis indicated that, relative to frogs maintained at 24°C, 2,392 genes were upregulated, whereas 2,031 genes were downregulated. Notably, genes associated with gluconeogenesis, such as *foxo1*, *g6pc1*, and *pck1*, exhibited significant upregulation, whereas genes related to glycolysis and glucose utilization, including *gck*, *pfkm*, and *ldhb*, were downregulated. Concurrently, cold exposure induced the expression of genes involved in fatty acid synthesis, desaturation, and cholesterol biosynthesis, such as *srebf1*, *acaca*, *fasn*, *scd*, *fads2*, *srebf2*, and *hmgcr*. Conversely, genes related to fatty acid β-oxidation, including *ppara*, *cpt1a*, *cpt1b*, *slc25a20*, *acadl*, *hadha*, and *hadhb*, were significantly suppressed. In alignment with this pattern, multiple components of the mitochondrial electron transport chain and ATP synthesis machinery were also downregulated. Additionally, gene groups involved in antioxidant defenses, such as *gpx4*, *gpx1*, *prdx2*, *prdx5*, *prdx6*, and ferritin-related genes, were upregulated. Representative transcriptomic changes were validated using quantitative PCR analysis.

**Conclusions:** Exposure to cold temperatures induced coordinated metabolic reprogramming in the liver of *X. laevis*. This reprogramming was characterized by the activation of gluconeogenesis, suppression of glycolysis, fatty acid β-oxidation, and oxidative phosphorylation, and induction of lipid and cholesterol biosynthetic pathways. These metabolic adjustments suggest that frogs acclimated to cold conditions adopt an energy-conserving metabolic strategy while sustaining glucose production and promoting lipid remodeling to adapt to low-temperature environments. These findings offer novel insights into the molecular mechanisms underlying cold adaptation in ectothermic vertebrates and underscore the pivotal role of the liver in managing transcriptional responses to cold stress.

## Background

Ambient temperature is a critical environmental factor that influences the physiological processes of ectothermic organisms, particularly in terms of metabolic regulation and cellular homeostasis. Exposure to low temperatures imposes energetic constraints that necessitate the reorganization of metabolic pathways to maintain homeostasis. Amphibians are valuable models for investigating temperature-dependent physiological responses, such as cold tolerance and metabolic adaptation. Freeze-tolerant species, such as *Rana sylvatica*, rapidly mobilize glucose from hepatic glycogen at the onset of freezing to accumulate cryoprotectants [1, 2]. These species can endure prolonged freezing periods through extensive metabolic reorganization and suppression of ATP-consuming processes during freezing [3–5]. These adaptations involve coordinated changes in carbohydrate metabolism, antioxidant defenses, and cellular stress responses, as demonstrated by physiological and transcriptomic analyses [3, 6]. Glucose metabolic responses to low temperature exposure have also been documented in amphibians, such as *X. laevis* [7]. Although this response may aid in maintaining energy homeostasis and cellular protection in cold environments, its molecular mechanisms remain unclear. Specifically, the transcriptional regulation that occurs in the liver during cold exposure and the regulation of glucose metabolism via gluconeogenesis and glycolysis are not fully understood. Recent transcriptomic analyses of the liver in freeze-tolerant amphibians have revealed differential expression of numerous cold-responsive genes [8]; however, the transcriptional responses in the livers of non-freeze-tolerant amphibians have not yet been fully elucidated. The liver, as a central metabolic organ, maintains systemic glucose homeostasis by regulating gluconeogenesis and glycolysis. Elucidating its transcriptional response is expected to provide key insights into the mechanisms underlying cold adaptation in amphibians. In addition to carbohydrate metabolism, lipid metabolism undergoes substantial changes in cold environments. Specifically, the synthesis of unsaturated fatty acids and alterations in fatty acid composition are essential for maintaining membrane fluidity and metabolic regulation and are considered the key mechanisms underlying cold adaptation in ectotherms [9, 10].

Adaptation to cold environments involves the regulation of membrane lipid composition and the implementation of energy-saving strategies through metabolic suppression. In hibernating mammals, including the Syrian hamster and 13-lined ground squirrel, marked reductions in metabolic rate and body temperature during torpor are accompanied by the suppression of energy-intensive processes, including transcription, translation, and protein synthesis [11, 12]. This enables survival during prolonged cold exposure and extended periods of fasting [13, 14]. This metabolic reorganization involves alterations in energy substrate utilization, modulation of mitochondrial function, and coordinated changes in protein degradation and proteostasis regulation, all of which contribute to efficient energy conservation. Enhanced defense mechanisms against oxidative stress have been reported in hibernators and cold-tolerant animals. During hibernation and arousal, antioxidant defenses are upregulated, mitigating oxidative damage induced by reoxygenation and rewarming [15, 16]. Recent studies suggest that lipid peroxidation suppression mechanisms involving *Gpx4* and ferritin contribute to cellular protection against cold stress and ferroptosis [3, 17]. Thus, cold adaptation is understood as an integrated strategy that includes not only the maintenance of membrane function through lipid remodeling but also the suppression of energy metabolism and the activation of cellular protection mechanisms. However, in non-hibernating amphibians, the changes in metabolic regulatory mechanisms and cellular protection mechanisms-related transcriptional programs in response to cold exposure remain largely unknown. Therefore, this study analyzed the hepatic transcriptome of *X. laevis* after exposure to 5 °C for 5 days using RNA-seq. The aim of this study was to elucidate the molecular basis of metabolic reprogramming in response to extreme thermal environments and advance the understanding of cold adaptation mechanisms in amphibians.

## Methods

### Animals and experimental design

In this study, adult African clawed frogs (*X. laevis*) were used as experimental subjects. Non-inbred wild-type frogs were used for the temporal analysis of serum parameters, whereas inbred wild-type frogs (J-strain) [18] were selected for RNA-seq and quantitative PCR analysis. All specimens were procured from the Amphibian Research Center at Hiroshima University (RRID: SCR_019015) through the National BioResource Project (Japan). The frogs were maintained at 24°C under a 12-hour light-dark cycle under standard housing conditions. For the cold exposure experiment, frogs were individually housed in separate 1-liter water containers and transferred, together with their housing water, to a 5°C incubator. In the time-course experiment, frogs were subjected to 5°C for up to 7 days. In the recovery experiment, frogs previously exposed to 5°C for 7 days were returned to 24°C for 2 days prior to sampling. For RNA-seq and qPCR analyses, frogs were exposed to 5°C for 5 days. At designated time points, the animals were euthanized. Blood samples were collected by cardiac puncture using a syringe and stored at −80°C. The liver tissue was promptly excised, flash-frozen in liquid nitrogen, and stored at −80°C until further analysis.

### Serum and hepatic metabolite measurements

Blood samples were centrifuged at 3,000 × g for 15 min at 4°C to facilitate serum separation. Serum glucose concentrations were determined using a GLUCOCARD G+ meter (Arkray, Kyoto, Japan). The NEFA C-test (FUJIFILM Wako Pure Chemical Corporation, Osaka, Japan) was used to measure the free fatty acid levels. Finally, the Triglyceride E-Test (FUJIFILM Wako Pure Chemical Corporation, Osaka, Japan) was used to measure triglyceride levels. To extract triglycerides from the liver, a previously reported protocol [19] was followed. The livers were homogenized in PBS, and a chloroform–methanol solution (2:1) was added to the homogenates. The samples were centrifuged at 18,000× g for 5 min at 4 °C, and the lower layer was collected and evaporated. The extracted lipids were dissolved in 100% isopropanol and hepatic triglyceride levels were measured using the Triglyceride E-Test (FUJIFILM Wako Pure Chemical Corporation, Osaka, Japan). Lipid peroxidation in liver tissue and serum was evaluated by measuring thiobarbituric acid reactive substances (TBARS) as previously described by Sugasawa et al. [19]. Briefly, liver tissues were homogenized in ice-cold buffer, and the resulting homogenates were used for TBARS analysis. Serum samples were analyzed directly. After reaction with thiobarbituric acid reagent, the absorbance at 532 nm was measured using a microplate reader. TBARS concentrations were calculated from a standard curve generated using standard solutions and expressed as nmol malondialdehyde equivalents. Lactate in serum was analyzed using a Beckman Coulter AU480 analyzer (Beckman Coulter, Krefeld, Germany) according to the manufacturer’s instructions.

### RNA extraction and sequencing

Total RNA was extracted from the liver tissue using TRIzol reagent (Thermo Fisher Scientific, USA) according to the manufacturer’s protocol. The RNA samples were treated with DNase I to eliminate genomic DNA contamination. Prior to library preparation, RNA quality was assessed, with RIN values ranging from 7.9–8.9.

RNA-seq libraries were constructed and sequenced by Novogene (Beijing, China) on an Illumina platform, producing 150 bp paired-end reads. A total of 278.3 million raw reads were obtained from the six libraries, with an average of 46.4 million reads per sample (range: 40.1–52.5 million). After filtering, 263.3 million clean reads remained, averaging 43.9 million reads per sample (range: 37.5–48.1 million). The Q20 and Q30 values exceeded 98% and 95%, respectively, indicating high-quality data.

### RNA-seq data processing and differential expression analysis

Raw sequence reads were filtered to remove adapter sequences and low-quality reads. Subsequently, high-quality reads were mapped to the *X. laevis* reference genome (GCF_017654675.1, Xenopus_laevis_version 10.1) using HISAT2 (version 2.0.5). Transcript assembly was performed using StringTie (version 1.3.3b), and gene expression levels were quantified using featureCounts (version 1.5.0-p3). Differential expression analysis was performed using DESeq2 (version 1.20.0). Genes with an adjusted *P* value (padj) < 0.05 and an absolute log₂ fold change (|log₂FC|) ≥ 1 were defined as differentially expressed genes (DEGs). Because *X. laevis* is an allotetraploid species, differential expression analyses were conducted separately for annotated L and S homoeologs. For readability, L/S suffixes were omitted in the main text unless distinction between homoeologs was relevant. Complete annotations are provided in Table S1.

### Functional enrichment analysis

Gene Ontology (GO) and Kyoto Encyclopedia of Genes and Genomes (KEGG) pathway enrichment analyses were executed utilizing clusterProfiler (version 3.8.1). Terms exhibiting padj < 0.05 were considered significantly enriched.

### Quantitative polymerase chain reaction (qPCR)

Total RNA was reverse transcribed into complementary DNA (cDNA) using the PrimeScript™ RT reagent kit (with gDNA Eraser, Takara Bio, Japan). qPCR was performed using the THUNDERBIRD™ Next SYBR® qPCR Mix (TOYOBO, Osaka, Japan) on a CFX Connect Real-Time PCR Detection System (Bio-Rad, Hercules, CA, USA). The PCR protocol included initial denaturation at 95°C for 60 s, followed by 39 cycles at 95°C for 10 s and 60°C for 30 s. Specific amplification was verified using melting curve analysis. Primer sequences are detailed in Table S2. Relative gene expression levels were determined using the 2^−ΔΔCt^ method and normalized to total eef1a1 expression detected using primers that recognize both L and S homoeologs.

### Statistical analysis

Data are presented as mean ± SEM. Statistical analyses were performed using one-way ANOVA followed by Tukey’s multiple comparisons test for the time-course serum glucose data (Fig. 1). Comparisons between the control (24°C) and cold-exposed (5°C for 5 days) groups in all other experiments were performed using the exact two-sided Mann–Whitney U test. Differences were considered statistically significant at *P* < 0.05.

**Fig. 1.**
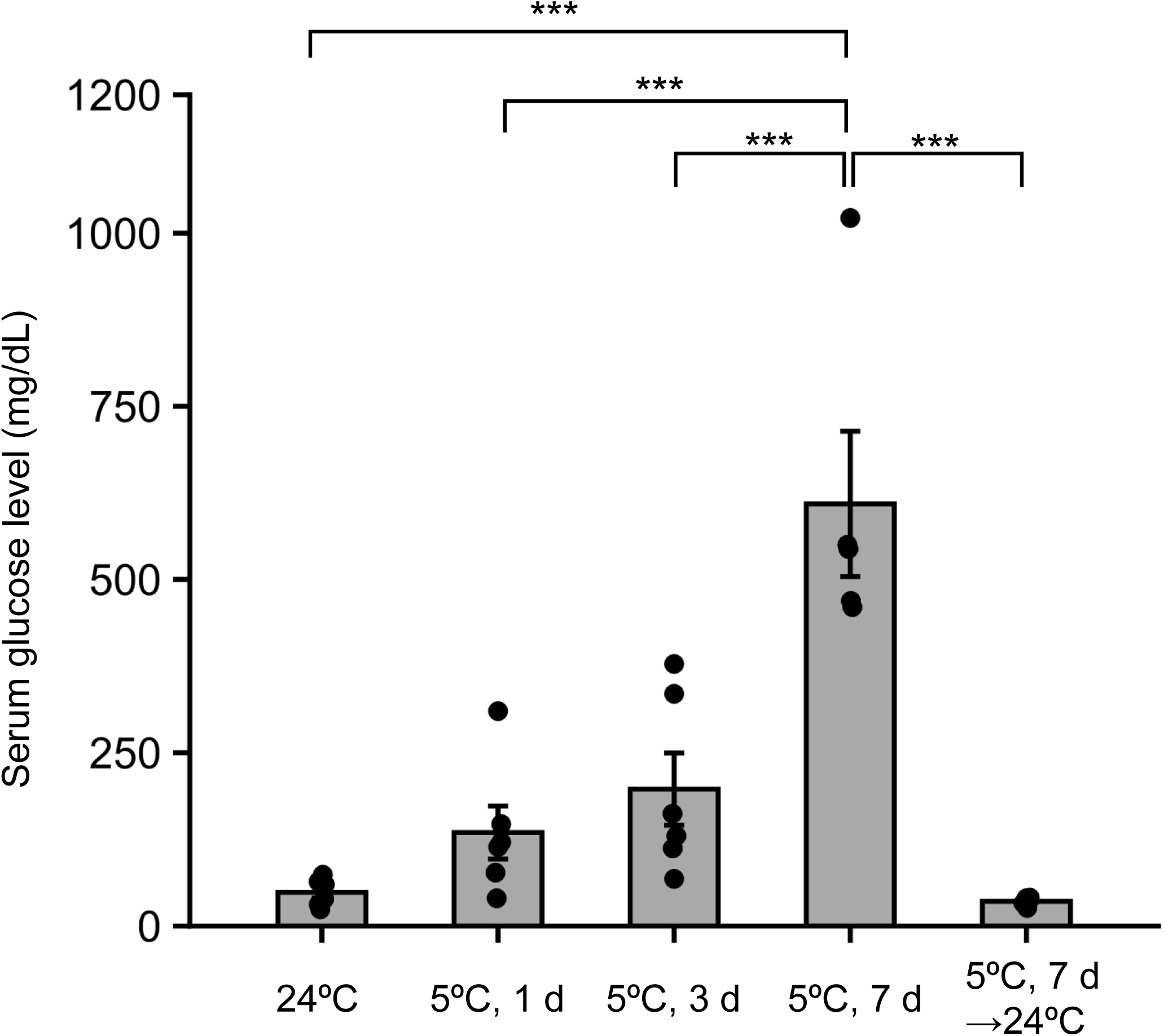
Serum glucose concentrations were measured in frogs maintained at 24°C (control) and in frogs exposed to 5°C for 1, 3, or 7 d, followed by rewarming at 24°C for 2 d. Data are presented as mean ± SEM (n = 5–10). Statistical significance was assessed using one-way ANOVA followed by Tukey’s multiple comparisons test. \*\*\**P* < 0.001.

## Results

### Cold exposure induces reversible hyperglycemia in *X. laevis*

To evaluate the physiological response to cold exposure, serum glucose concentrations were monitored over time in frogs maintained at 24°C and those exposed to 5°C. Serum glucose levels increased progressively during cold exposure and remained elevated throughout the exposure period, reaching significantly higher levels after 7 days compared to the control group (Fig. 1). Following 7 days of exposure to 5°C, the frogs were returned to 24°C for 2 days, after which serum glucose concentrations decreased to near-baseline levels (Fig. 1).

### Transcriptomic profiling reveals widespread gene expression changes during cold exposure

To investigate the molecular mechanisms underlying the sustained hyperglycemic response to cold exposure (Fig. 1), liver RNA-seq analysis was performed using samples collected after 5 days of exposure to 5°C. The differential expression analysis identified 4,423 genes exhibiting significant changes (padj < 0.05, |log₂ fold change| ≥ 1), comprising 2,392 upregulated and 2,031 downregulated genes in the livers of cold-exposed frogs (Fig. 2a). Hierarchical clustering analysis of the DEGs distinctly separated the control and cold-exposed groups, indicating substantial transcriptional alterations in response to cold exposure (Fig. 2b). Gene Ontology (GO) enrichment analysis demonstrated significant enrichment of terms related to transcription regulator activity, DNA-binding transcription factor activity, inhibitor activity, lipid binding, and iron ion binding among the DEGs (Fig. 2c). KEGG pathway analysis of the upregulated genes revealed significant enrichment in the FoxO signaling, PPAR signaling, fatty acid metabolism, ferroptosis, adipocytokine signaling, insulin signaling, and p53 signaling pathways (Fig. 2d). Conversely, downregulated genes were significantly enriched in pathways related to transcription and translation, including RNA polymerase, spliceosome, mRNA surveillance, aminoacyl-tRNA biosynthesis, and ribosome biogenesis (Fig. 2e). These enriched pathways suggest that cold exposure induces extensive transcriptional changes in hepatic metabolic regulation, stress responses, and protein synthesis, which were further examined in subsequent analyses of the study.

**Fig. 2.**
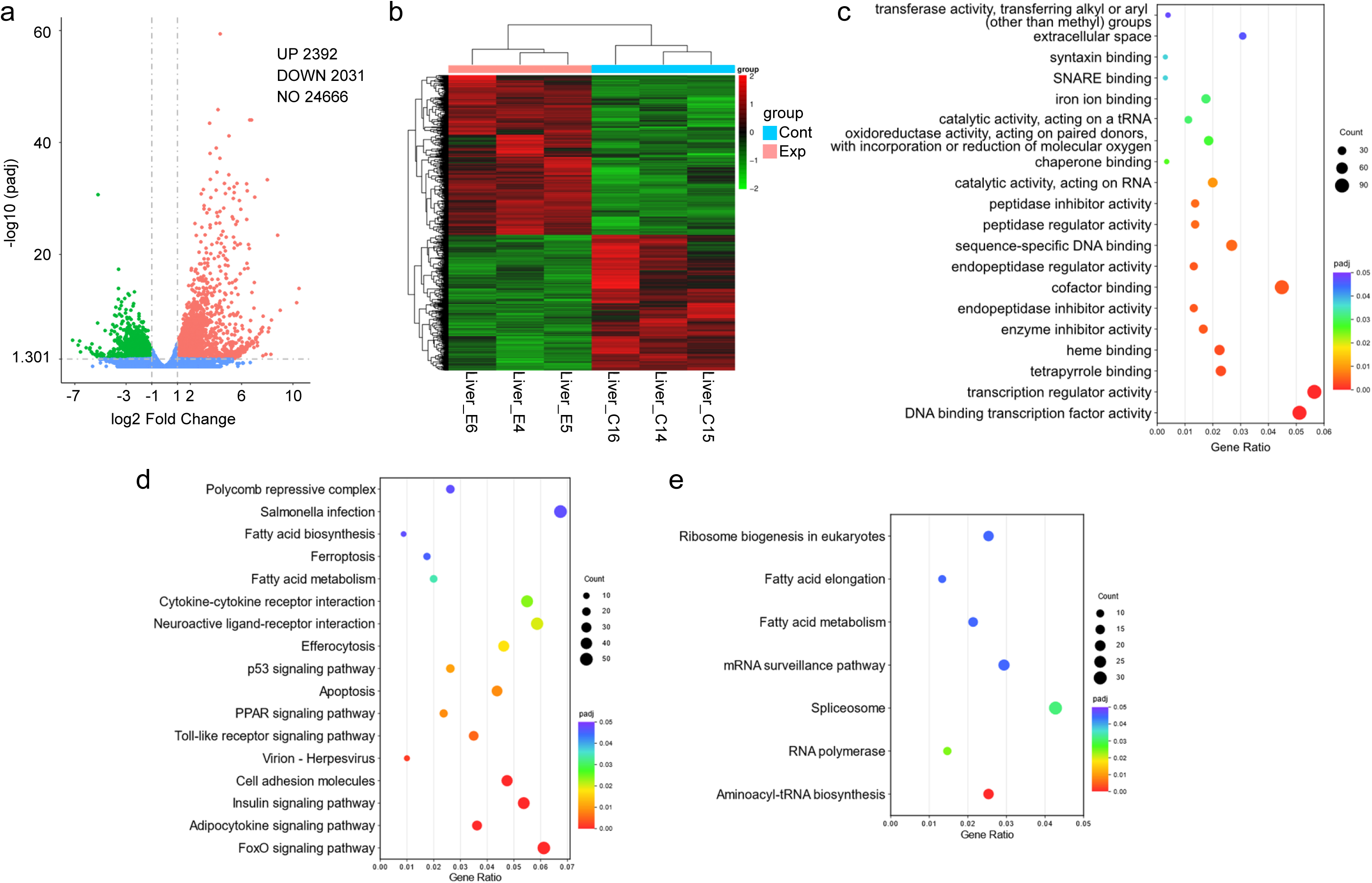
RNA-seq analysis was performed on liver samples from control frogs maintained at 24°C (Cont) and cold-exposed frogs (5°C, 5 d; Exp). (a) Volcano plot showing differentially expressed genes (DEGs). (b) Heatmap showing hierarchical clustering of DEGs. (c) Gene Ontology (GO) enrichment analysis of DEGs. (d) KEGG pathway enrichment analysis of upregulated genes. (e) KEGG pathway enrichment analysis of downregulated genes. DEGs were defined as genes with an adjusted *P* value (padj) < 0.05 and an absolute log₂ fold change ≥ 1. Dot size represents the number of genes associated with each term, and color indicates the padj value.

### Biochemical characterization of frogs following five days of cold exposure

To clarify the physiological state linked to transcriptomic changes, serum and hepatic biochemical parameters were evaluated in frogs after five days of cold exposure (Table 1). Serum glucose concentrations were significantly higher in cold-exposed frogs compared to the control group. In contrast, lactate concentrations were notably reduced, whereas serum non-esterified fatty acid (NEFA) and triglyceride concentrations remained unchanged. The hepatic triglyceride content did not differ significantly between the groups. Furthermore, the TBARS levels in both serum and liver were comparable between the control and cold-exposed frogs (Table 1).

**Table 1.** Serum and hepatic metabolite concentrations in control and cold-exposed frogs.

|  |  | Control (24°C) | Cold exposure (5°C, 5 d) |
| --- | --- | --- | --- |
| Serum | Glucose (mg/dL) | 37.5 ± 10.77 | 273.83 ± 24.27* |
|  | Triglyceride (mg/dL) | 254.85 ± 60.04 | 78.15 ± 7.94 |
|  | Free fatty acid (mEq/L) | 0.32 ± 0.03 | 0.64 ± 0.17 |
|  | Lactate (mg/dL) | 84.48 ± 16.24 | 13.64 ± 1.58* |
|  | TBARS (nmol/mL) | 3.90 ± 0.76 | 3.91 ± 0.49 |
| Liver | Triglyceride (mg/g tissue) | 9.89 ± 2.56 | 12.06 ± 2.27 |
|  | TBARS (nmol/mg tissue) | 0.063 ± 0.010 | 0.064 ± 0.004 |

Values are presented as mean ± SEM (n = 5–6 per group). Statistical significance between groups was determined using an exact two-sided Mann–Whitney U-test. \* $P < 0.05$ .

### Cold exposure is associated with transcriptional changes in FOXO1-related signaling and hepatic glucose metabolism

To elucidate the molecular mechanisms underlying cold-induced hyperglycemia, we analyzed the genes implicated in FOXO1 signaling and hepatic glucose metabolism. FOXO1 is the principal transcriptional regulator of gluconeogenic gene expression. RNA-seq demonstrated an upregulation of *foxo1* and its coactivator *prmt1* in the livers of frogs exposed to cold conditions (Table 2 and Fig. 3), indicating the activation of FOXO1-associated transcriptional programs. Corresponding to the enhanced gluconeogenic capacity, the expression levels of the gluconeogenic genes *g6pc* and *pck1* were significantly elevated in cold-exposed frogs (Table 2 and Fig. 3). Conversely, the expression of several genes involved in glucose utilization was reduced. The glycolytic genes *gck* and *pfkm* were downregulated, suggesting suppression of glycolytic flux (Table 2 and Fig. 3). Additionally, the expression of *ldhb*, which preferentially catalyzes the conversion of lactate to pyruvate, decreased (Table 2 and Fig. 3). The expression of *pc*, which connects pyruvate metabolism to the tricarboxylic acid cycle and gluconeogenesis, was also downregulated (Table 2 and Fig. 3). The RNA-seq results were corroborated by qPCR analysis of selected genes, including *foxo1*, *g6pc1*, and *pck1*, whereas *prmt1* showed a similar trend that did not reach statistical significance (*P* = 0.06) (Fig. 6a). Furthermore, the expression of *gck*, a critical enzyme involved in hepatic glucose utilization, was significantly reduced (Fig. 6a).

**Fig. 3.**
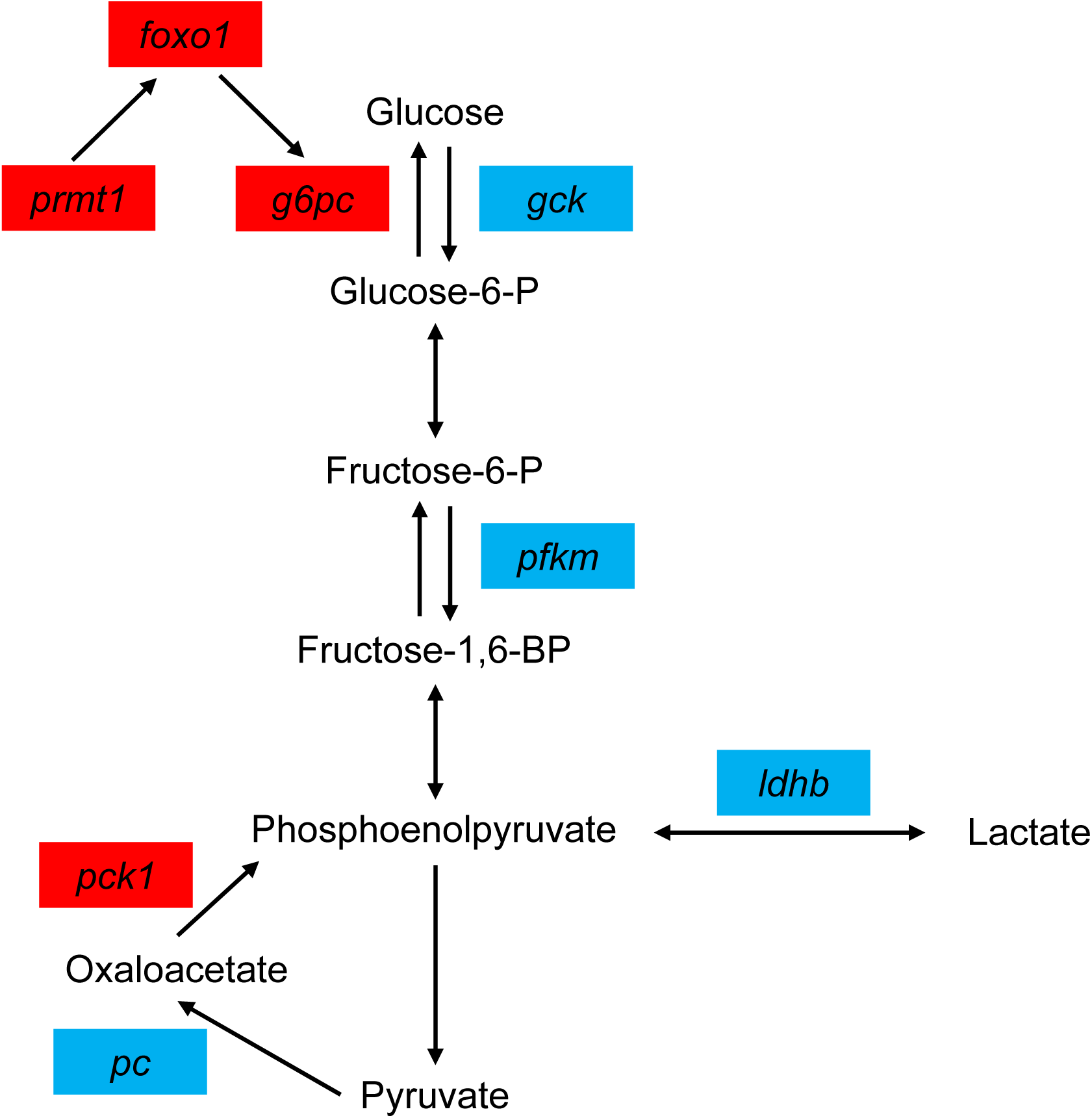
Schematic representation of transcriptional changes in hepatic gluconeogenesis and glycolysis pathways based on RNA-seq analysis. Genes significantly upregulated by cold exposure are shown in red, whereas significantly downregulated genes are shown in blue. Metabolic intermediates are shown in black. Arrows indicate metabolic conversions or regulatory relationships, and bidirectional arrows indicate the reversible reactions.

**Table 2.** Differential expression of genes involved in gluconeogenesis and glycolysis.

| Gene (Xenbase) | Pathway | Homoeologs | log<sub>2</sub>FC | padj |
| --- | --- | --- | --- | --- |
| ***foxo1*** | **FOXO signaling** | L | +3.17 | 7.9.E-07 |
|  |  | S | +2.70 | 9.9.E-05 |
| ***prmt1*** | **FOXO1 regulation (post-translational modification)** | L | +1.29 | 5.7.E-03 |
| ***g6pc1*** | **Gluconeogenesis** | L | +5.41 | 8.5.E-33 |
|  |  | S | +5.04 | 1.4.E-15 |
| ***g6pc1.2*** | **Gluconeogenesis** | S | +1.85 | 1.3.E-02 |
| ***g6pc1.3*** | **Gluconeogenesis** | L | +6.09 | 2.0.E-07 |
|  |  | S | +6.44 | 4.4.E-32 |
| ***pck1*** | **Gluconeogenesis** | L | +2.40 | 1.2.E-04 |
| ***pc*** | **Gluconeogenesis** | L | -1.43 | 4.1.E-02 |
| ***gck*** | **Glycolysis / Glucose utilization** | L | -2.34 | 1.1.E-03 |
|  |  | S | -1.84 | 1.1.E-03 |
| ***pfkm*** | **Glycolysis** | S | -1.43 | 9.8.E-03 |
| ***ldhb*** | **Glycolysis** | L | -1.98 | 9.5.E-04 |
|  |  | S | -1.76 | 3.4.E-02 |

RNA-seq analysis showing expression changes in genes involved in gluconeogenesis and glycolysis in the liver of *X. laevis* exposed to 5°C for 5 d. Gene names follow Xenbase annotation. Homoeologs indicate gene copies derived from the L and S subgenomes. The log<sub>2</sub> fold change (log<sub>2</sub>FC) represents expression changes relative to the control group maintained at 24°C, and adjusted *P* values (padj) indicate statistical significance.

### Cold exposure alters the transcriptional regulation of lipid and cholesterol metabolic pathways

We investigated the transcriptional alterations in lipogenesis and cholesterol biosynthesis following cold exposure. RNA-seq demonstrated an upregulation of *srebf1*, a principal transcriptional regulator of lipogenesis, along with increased expression of its downstream target genes, *acaca* and *fasn* (Table 3 and Fig. 4a). Additionally, the expression of genes involved in fatty acid elongation and desaturation, including *elovl6*, *elovl5*, *scd*, and *fads2*, was significantly increased, indicating an enhanced capacity for fatty acid remodeling and unsaturated fatty acid synthesis (Table 3 and Fig. 4a). Genes involved in cholesterol biosynthesis were also upregulated in response to cold exposure. The expression of *srebf2*, a critical transcriptional regulator of cholesterol metabolism, was increased, accompanied by elevated expression of its downstream target genes involved in cholesterol biosynthesis, including *hmgcr*, *fdft1*, *sqle*, and *cyp51a1* (Table 3 and Fig. 4b). The RNA-seq results were generally consistent with qPCR analysis of selected genes. Significant differences were confirmed for *srebf1*, *acaca*, *fasn*, *scd*, *fads2*, *srebf2*, and *hmgcr*, while *cyp51a1* exhibited a similar trend that did not reach statistical significance (Fig. 6b, c).

**Fig. 4.**
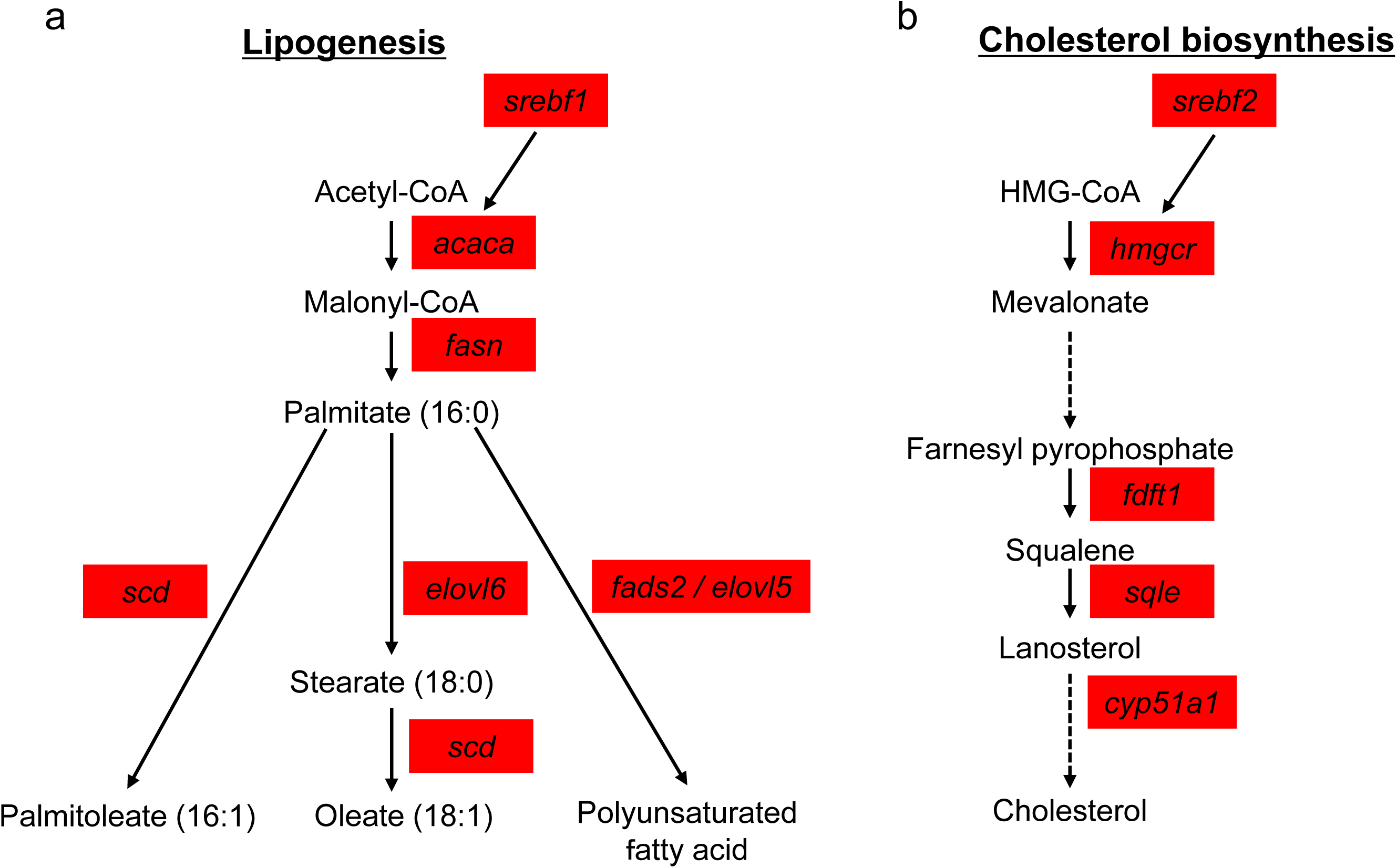
Schematic representation of transcriptional changes in genes involved in hepatic lipogenesis (a) and cholesterol biosynthesis (b) based on RNA-seq analysis. Genes significantly upregulated by cold exposure are shown in red. Metabolic intermediates are indicated in black. Arrows indicate metabolic conversions or regulatory relationships, and bidirectional arrows indicate reversible reactions. Dashed arrows indicate pathways involving multiple intermediate

**Table 3.**
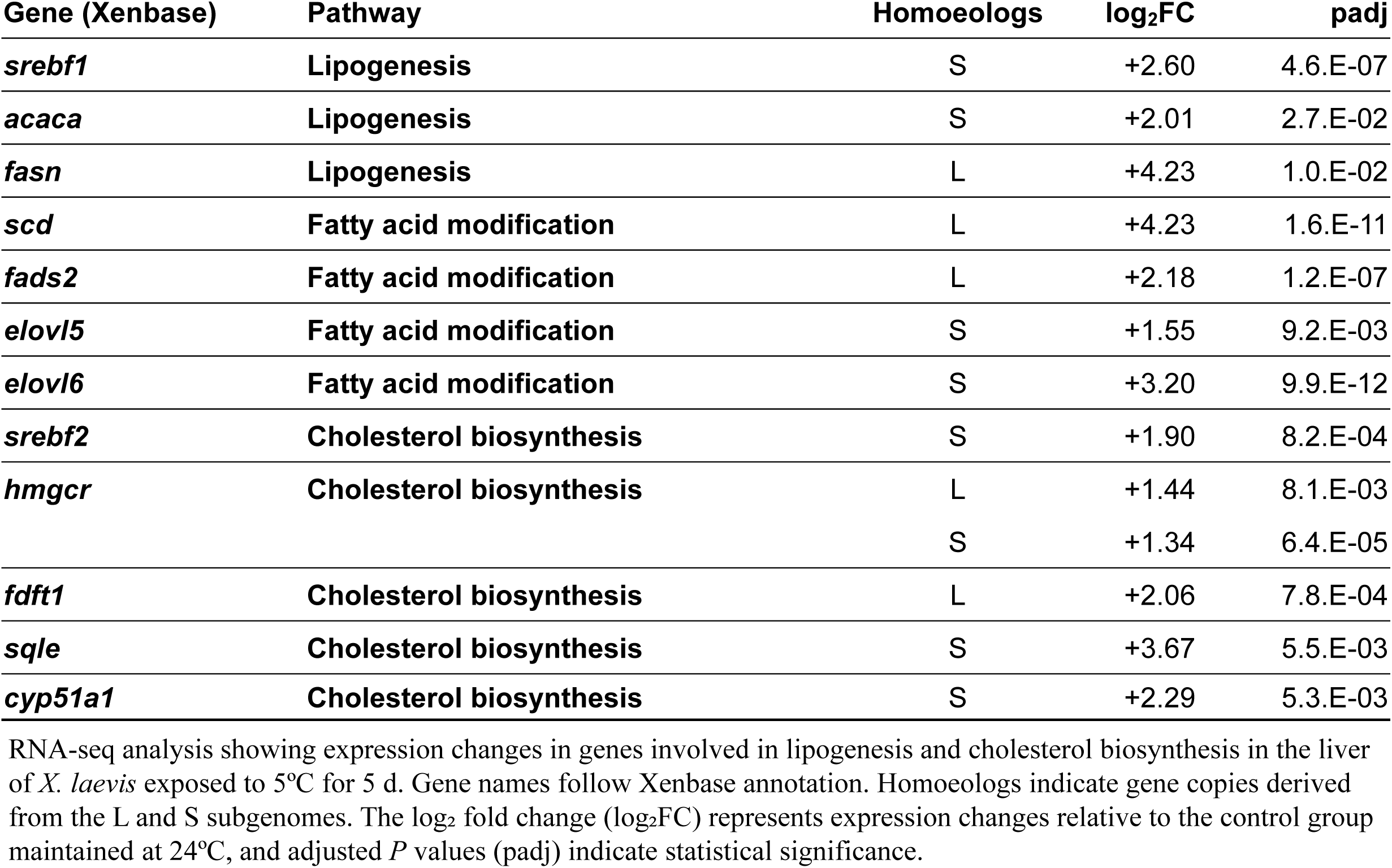
Differential expression of genes involved in lipogenesis and cholesterol biosynthesis.

### Cold exposure induces transcriptional changes in fatty acid β-oxidation, electron transport chain, and antioxidant defense pathways

To examine the impact of cold exposure on hepatic energy metabolism and oxidative stress responses, we analyzed genes associated with fatty acid β-oxidation, the electron transport chain, and antioxidant defense pathways. RNA-seq indicated a significant downregulation of *ppara*, a principal regulator of fatty acid catabolism, along with decreased expression of genes involved in mitochondrial fatty acid transport and β-oxidation, such as *cpt1a*, *cpt1b*, *slc25a20*, *acadl*, *hadha*, *hadhb*, and *acaa2* (Table 4 and Fig. 5a). In addition, genes encoding components of the mitochondrial electron transport chain were downregulated. Specifically, reduced expression was noted for genes linked to Complex I (*mt-nd1*, *mt-nd4*, *mt-nd5*, *ndufs1*, and *ndufv2*), Complex II (*sdhd*), Complex III (*cyc1*, *uqcrb*, and *uqcrq*), the electron carrier *cycs*, Complex IV (*cox5b*, *cox6c*, *cox7a2*, and *cox7b*), and Complex V (*atp5f1b*, *atp5f1d*, and *atp5pb*) (Table 4 and Fig. 5b).

**Fig. 5.**
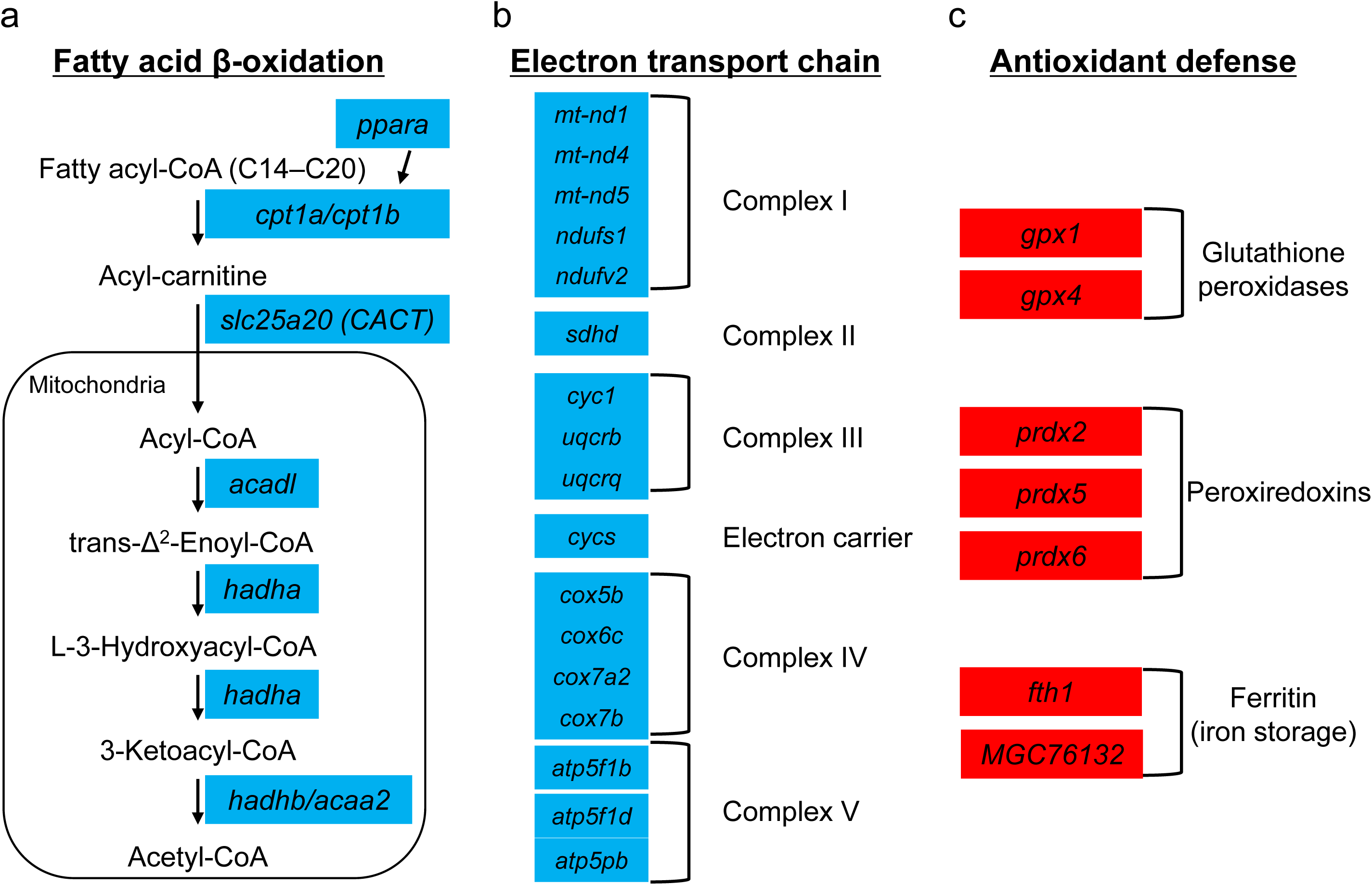
Schematic representation of transcriptional changes in genes involved in hepatic fatty acid β-oxidation (a), electron transport chain (b), and antioxidant defense (c) based on RNA-seq analysis. Genes significantly upregulated and downregulated by cold exposure are shown in red and blue, respectively. Arrows indicate metabolic conversions or regulatory relationships.

**Table 4.**
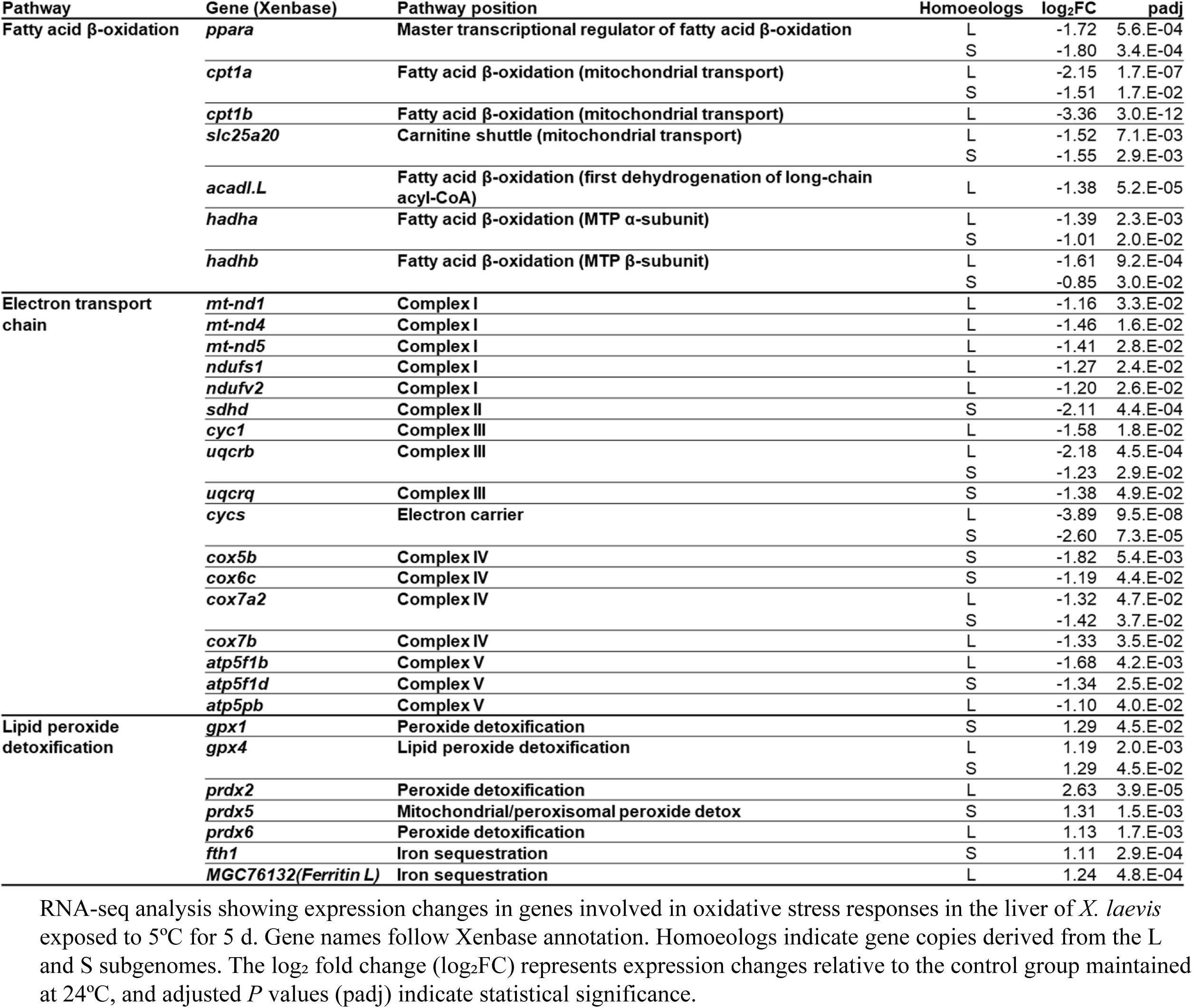
Differential expression of genes involved in oxidative stress responses.

Conversely, several antioxidant defense genes were upregulated in response to exposure to cold. There was a significant increase in the expression of glutathione peroxidase genes (*gpx1* and *gpx4*), peroxiredoxin genes (*prdx2*, *prdx5*, and *prdx6*), and ferritin-related genes (*fth1* and *MGC76132*) (Table 4 and Fig. 5c). The RNA-seq results were further examined using qPCR analysis. Significant decreases in *ppara*, *cpt1a*, and *cpt1b* expression and an increase in *gpx4* expression were confirmed in cold-exposed frogs (Fig. 6d).

**Fig. 6.**
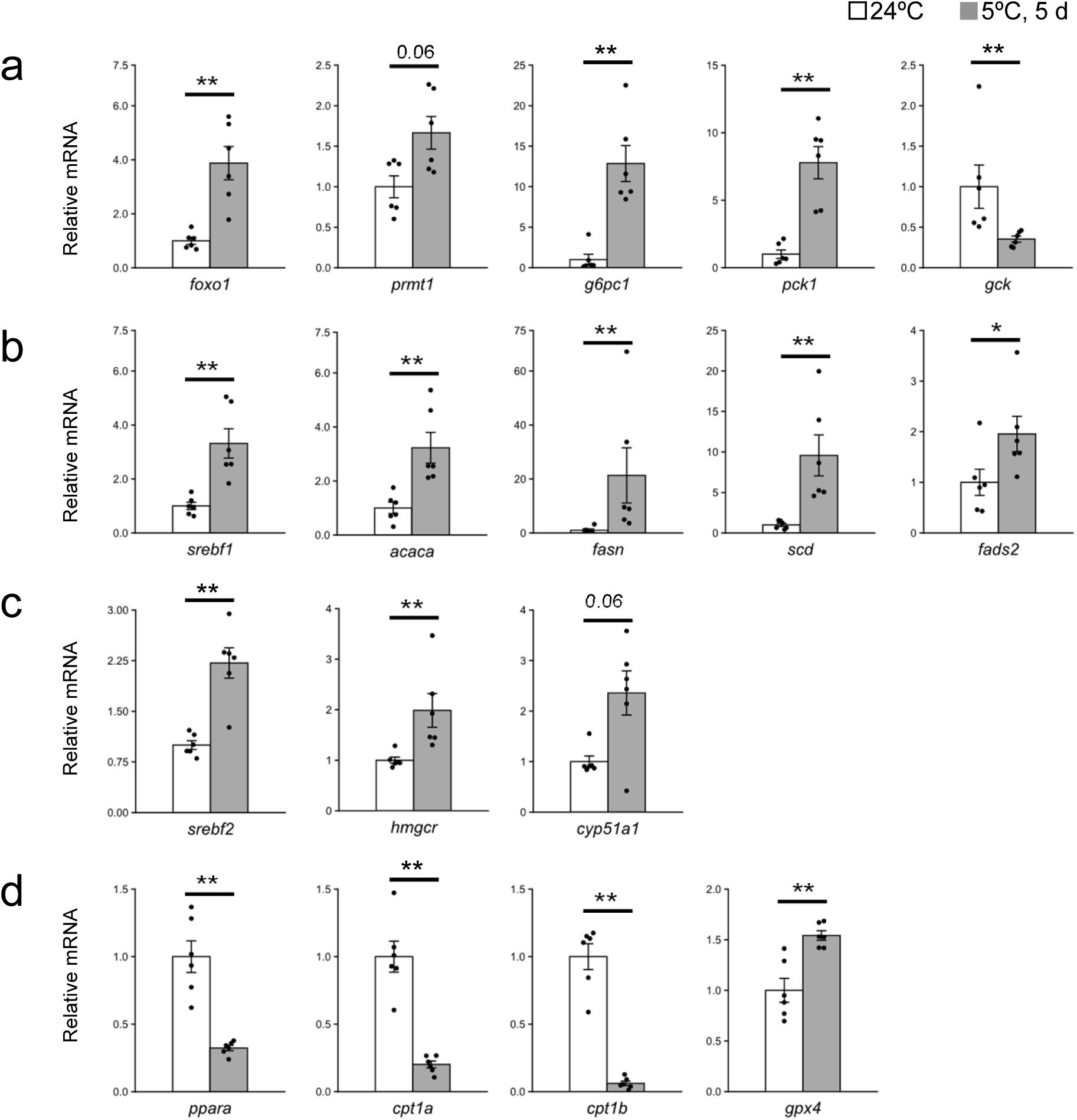
Relative mRNA expression levels of genes involved in gluconeogenesis and glycolysis (a), lipid biosynthesis (b), cholesterol biosynthesis (c), and fatty acid β-oxidation and antioxidant defense (d) in the livers of *X. laevis* maintained at 24°C or exposed to 5°C for 5 d (n = 6 per group). Data are presented as individual data points with the mean + SEM. Statistical significance was assessed using an exact two-sided Mann–Whitney U-test. \**P*<0.05, \*\**P* < 0.01. Exact *P* values are shown above the bars for comparisons that did not reach statistical significance.

### Cold exposure alters transcription of protein turnover pathways

Finally, we examined the transcriptional alterations in genes implicated in protein metabolism. Genes associated with aminoacyl-tRNA synthetases were predominantly downregulated, indicating reduced translational activity (Table 5). Conversely, the expression of genes linked to the ubiquitin–proteasome system, including E3 ubiquitin ligase *fbxo32* and ubiquitin-conjugating enzymes (*ube2d1*, *ube2r2*, *ube2s*, and *ube2z*), was upregulated (Table 5). Furthermore, the autophagy-related gene *map1lc3b* exhibited increased expression (Table 5).

**Table 5.** Differential expression of genes involved in protein turnover pathways.

| Category | Gene (Xenbase) | Homoeologs | log<sub>2</sub>FC | padj | Direction |
| --- | --- | --- | --- | --- | --- |
| Aminoacyl-tRNA synthetase | *dars2* | L | -2.96 | 8.2.E-07 | ↓ |
|  | *lars1* | L | -1.24 | 1.0.E-03 | ↓ |
|  | *mars1* | L | -1.55 | 2.6.E-03 | ↓ |
|  | *mars2* | L | -3.45 | 2.5.E-04 | ↓ |
|  | *farsa* | L | -1.59 | 1.3.E-03 | ↓ |
|  |  | S | -1.90 | 2.4.E-04 | ↓ |
|  | *pars2* | L | -1.72 | 2.5.E-02 | ↓ |
|  | *sars2* | L | -1.68 | 4.1.E-02 | ↓ |
|  | *tars1* | L | -1.31 | 1.4.E-02 | ↓ |
|  | *wars1* | L | -1.81 | 2.0.E-03 | ↓ |
|  |  | S | -1.10 | 2.6.E-02 | ↓ |
|  | *wars2* | S | -1.97 | 3.6.E-02 | ↓ |
|  | *yars1* | L | -2.21 | 4.1.E-05 | ↓ |
|  | *vars1* | S | -1.25 | 4.6.E-02 | ↓ |
|  | *vars2* | L | -1.38 | 4.6.E-03 | ↓ |
| Ubiquitin–proteasome system | *fbxo32* | L | 4.16 | 4.1.E-11 | ↑ |
|  |  | S | 4.88 | 1.0.E-19 | ↑ |
|  | *ube2d1* | L | 1.22 | 6.7.E-04 | ↑ |
|  | *ube2r2* | S | 2.11 | 1.1.E-03 | ↑ |
|  | *ube2s* | L | 1.31 | 1.2.E-03 | ↑ |
|  | *ube2z* | S | 1.34 | 2.0.E-04 | ↑ |
| Autophagy | *map1lc3b* | L | 1.15 | 5.6.E-03 | ↑ |
|  |  | S | 1.39 | 1.1.E-02 | ↑ |

Genes involved in aminoacyl-tRNA biosynthesis, the ubiquitin–proteasome system (UPS), and autophagy are shown. Gene names follow Xenbase annotation. Homoeologs indicate gene copies derived from the L and S subgenomes. Log<sub>2</sub> fold change (log<sub>2</sub>FC) represents expression changes relative to the control group (24°C). Adjusted *P* values (padj) indicate statistical significance. Arrows indicate the direction of differential expression (↑, upregulated; ↓, downregulated) in cold-exposed frogs (5°C, 5 d) relative to controls.

## Discussion

In this study, we examined the hepatic transcriptional response of *X. laevis* to cold exposure using RNA-seq and qPCR analyses. Cold exposure resulted in significant hyperglycemia and extensive transcriptional reprogramming in the liver. This reprogramming was characterized by the activation of the FOXO1-associated gluconeogenic pathway, induction of fatty acid and cholesterol biosynthetic pathways, suppression of fatty acid β-oxidation and oxidative phosphorylation, enhancement of antioxidant defense mechanisms, and downregulation of protein synthesis pathways. These findings indicate that the liver undergoes a coordinated metabolic reorganization to maintain energy homeostasis under cold stress. A prominent response to cold exposure was the activation of the FOXO1 signaling pathway and hepatic gluconeogenesis. The expression of *foxo1*, *g6pc1*, and *pck1* was significantly upregulated, whereas that of *gck* was decreased. Although not statistically significant, *prmt1* expression also tended to increase (*P* = 0.06), further supporting activation of the FOXO1-associated gluconeogenic pathway. These transcriptional changes were accompanied by a marked increase in circulating blood glucose levels. The significant upregulation of the gluconeogenic genes *g6pc1* and *pck1*, consistent with the upregulation of *foxo1* expression, suggests that the FOXO1-associated gluconeogenic program is activated in the liver under cold exposure.

Acute cold exposure induces profound metabolic remodeling in mammalian liver. FOXO1 is a key transcriptional regulator of hepatic glucose production through the activation of gluconeogenic genes, such as G6pc and Pck1 [20, 21]. Additionally, recent studies on freeze-tolerant wood frogs have demonstrated the activation of FOXO3 signaling during freezing and thawing, suggesting a role for FOXO-mediated regulation in coordinating antioxidant defenses and metabolic adaptation to freezing stress [22]. Combined with the findings of the present study, this observation suggests that FOXO signaling activation may be a conserved adaptive response to cold stress in vertebrates. However, whether FOXO1 is directly involved in the transcriptional response observed in the livers of cold-exposed *X. laevis* remains to be elucidated. Further studies examining FOXO1 protein levels, phosphorylation status, and nuclear localization will be necessary to confirm its functional involvement.

Following cold exposure, serum lactate levels decreased notably. Lactate, the primary end product of glycolysis, is a prevalent marker of glycolytic activity. Consequently, the reduction in circulating lactate, coupled with the observed downregulation of glycolysis-related genes such as *gck*, *pfkm*, and *ldhb*, indicates the suppression of glycolytic metabolism under cold conditions. A comparable decrease in lactate levels has been reported in hibernating ground squirrels during torpor, consistent with the profound metabolic suppression characteristic of hibernation[23]. Reduced glycolytic activity and limited lactate accumulation during torpor have been reported in hibernating ground squirrels, reflecting a shift away from glucose catabolism and contributing to metabolic suppression [24]. Conversely, in freeze-tolerant frogs, glycolytic activity and lactate accumulation increase upon freezing, illustrating a distinct metabolic strategy that facilitates freeze tolerance [25]. These observations suggest that vertebrates employ various metabolic strategies to adapt to cold stress conditions. While freeze-tolerant frogs enhance glycolytic flux and accumulate lactate during freezing, *X. laevis* demonstrated a reduction in circulating lactate concentration and transcriptional suppression of glycolytic genes. This response aligns more closely with a metabolic program characterized by decreased glycolytic activity and overall energy conservation, rather than the activation of anaerobic metabolism. Exposure to cold temperatures elicited significant transcriptional alterations in lipid metabolism. Genes associated with fatty acid synthesis and unsaturation, such as *srebf1*, *acaca*, *fasn*, *scd*, *fads2*, *elovl5*, and *elovl6*, were upregulated, indicating enhanced lipid synthesis and fatty acid remodeling. Simultaneously, genes involved in cholesterol biosynthesis, including *srebf2*, *hmgcr*, *fdft1*, *sqle*, and *cyp51a1* were activated. Notably, despite activating the lipid synthesis pathway, hepatic triglyceride levels remained constant, suggesting that the transcriptional response may not primarily facilitate lipid storage. In addition to the upregulation of genes related to fatty acid biosynthesis, the expression of desaturases *scd* and *fads2* and elongases *elovl5* and *elovl6* increased in response to cold exposure. These enzymes are pivotal in determining the unsaturation and carbon chain length of membrane lipids, and their induction may contribute to membrane lipid remodeling during cold adaptation. These changes are consistent with the concept of homeoviscous adaptation, whereby ectotherms remodel their membrane lipid composition through the coordinated regulation of fatty acid desaturation and cholesterol metabolism to maintain membrane fluidity under cold conditions [10, 26, 27]. In contrast to the induction of the lipid biosynthesis pathway, the expression of genes involved in mitochondrial fatty acid β-oxidation was generally downregulated. The downregulation of *ppara*, *cpt1a*, *cpt1b*, *slc25a20*, *acadl*, *hadha*, *hadhb*, and *acaa2* indicates a reduction in mitochondrial fatty acid utilization under cold exposure. Consistent with this observation, multiple genes constituting the electron transport chain and oxidative phosphorylation pathway were downregulated. These findings suggest that mitochondrial oxidative metabolism and ATP production capacity are diminished in frogs exposed to cold temperatures. Such suppression of mitochondrial energy metabolism may represent an energy conservation strategy under conditions of reduced metabolic demand and is consistent with the characteristic metabolic rate depression commonly observed in ectotherms and hibernating vertebrates [11, 28]. In parallel with the suppression of mitochondrial oxidative metabolism, several antioxidant-related genes, including *gpx1*, *gpx4*, *prdx2*, *prdx5*, and *prdx6*, as well as ferritin-associated genes, were upregulated. Notably, TBARS concentrations in both serum and liver remained unchanged despite the upregulation of antioxidant-related genes, suggesting that the induction of antioxidant defense pathways may contribute to preserving redox homeostasis during cold exposure. The cold-induced transcriptomic response was characterized by the significant downregulation of pathways associated with RNA processing and protein synthesis, such as RNA polymerase, spliceosome, ribosome biogenesis, and aminoacyl-tRNA biosynthesis. Given that protein synthesis is among the most energy-intensive cellular processes, the suppression of translational capacity may represent an important energy-conservation strategy during cold exposure, consistent with the metabolic rate depression observed in ectotherms and hibernating vertebrates [11, 14]. Along with diminished fatty acid β-oxidation and oxidative phosphorylation, these observations suggest the presence of a comprehensive energy-saving mechanism in the cold-exposed frog liver.

Cold exposure results in extensive metabolic reprogramming in the *X. laevis* liver, as evidenced by transcriptional responses that include enhanced gluconeogenesis, activation of lipid and cholesterol biosynthetic pathways, suppression of mitochondrial energy metabolism, reinforcement of antioxidant defense systems, and reduction in protein synthesis. These coordinated changes likely constitute an adaptive strategy to maintain glucose homeostasis while minimizing energy expenditure during cold stress. Our findings collectively support a model in which cold exposure triggers dual metabolic reprogramming in the *X. laevis* liver. Activation of the FOXO1-related gluconeogenic pathway aids in maintaining circulating glucose levels, whereas inhibition of fatty acid β-oxidation, oxidative phosphorylation, and protein synthesis indicates the initiation of an energy conservation program. Concurrently, the induction of fatty acid synthesis, desaturation, and cholesterol biosynthetic pathways may facilitate lipid remodeling and adaptation to low-temperature environments. In contrast to freeze-tolerant frogs, which accumulate significantly elevated levels of glucose as a cryoprotectant, *X. laevis* exhibits a more moderate increase in circulating glucose. This finding indicates that increased glucose production may play a role in maintaining energy homeostasis during cold exposure, independent of the freeze protection mechanisms. Although this study offers extensive transcriptomic insights into cold-induced metabolic reprogramming, the functional implications of these transcriptional changes remain to be confirmed. Future research should focus on assessing metabolite flux, membrane lipid composition, and mitochondrial function to substantiate these transcriptomic predictions.

## Conclusions

In summary, cold exposure induced significant transcriptional reprogramming in the liver of *X. laevis*, accompanied by pronounced hyperglycemia and coordinated alterations across various metabolic pathways. The responses include activation of the FOXO1-associated gluconeogenic program and suppression of glycolysis, fatty acid β-oxidation, oxidative phosphorylation, and protein synthesis. These findings support a model in which cold exposure initiates dual metabolic reprogramming in the liver. Activation of gluconeogenesis aids in maintaining circulating glucose levels, whereas suppression of energy-consuming processes establishes an energy-sparing program that reduces metabolic demand. Concurrently, the induction of lipid and cholesterol biosynthetic pathways likely facilitates lipid remodeling, aiding cellular adaptation to low temperatures.

Collectively, our results indicate that the liver plays a central and coordinated role in the systemic metabolic adaptation to cold stress in *X. laevis* by integrating pathways of glucose homeostasis, energy conservation, and lipid remodeling (Fig. 7). This study offers new insights into the molecular mechanisms underlying adaptation to extreme thermal environments, including cold, in ectothermic vertebrates, and establishes a foundation for future research into the regulatory networks governing temperature-dependent metabolic plasticity.

**Fig. 7.**
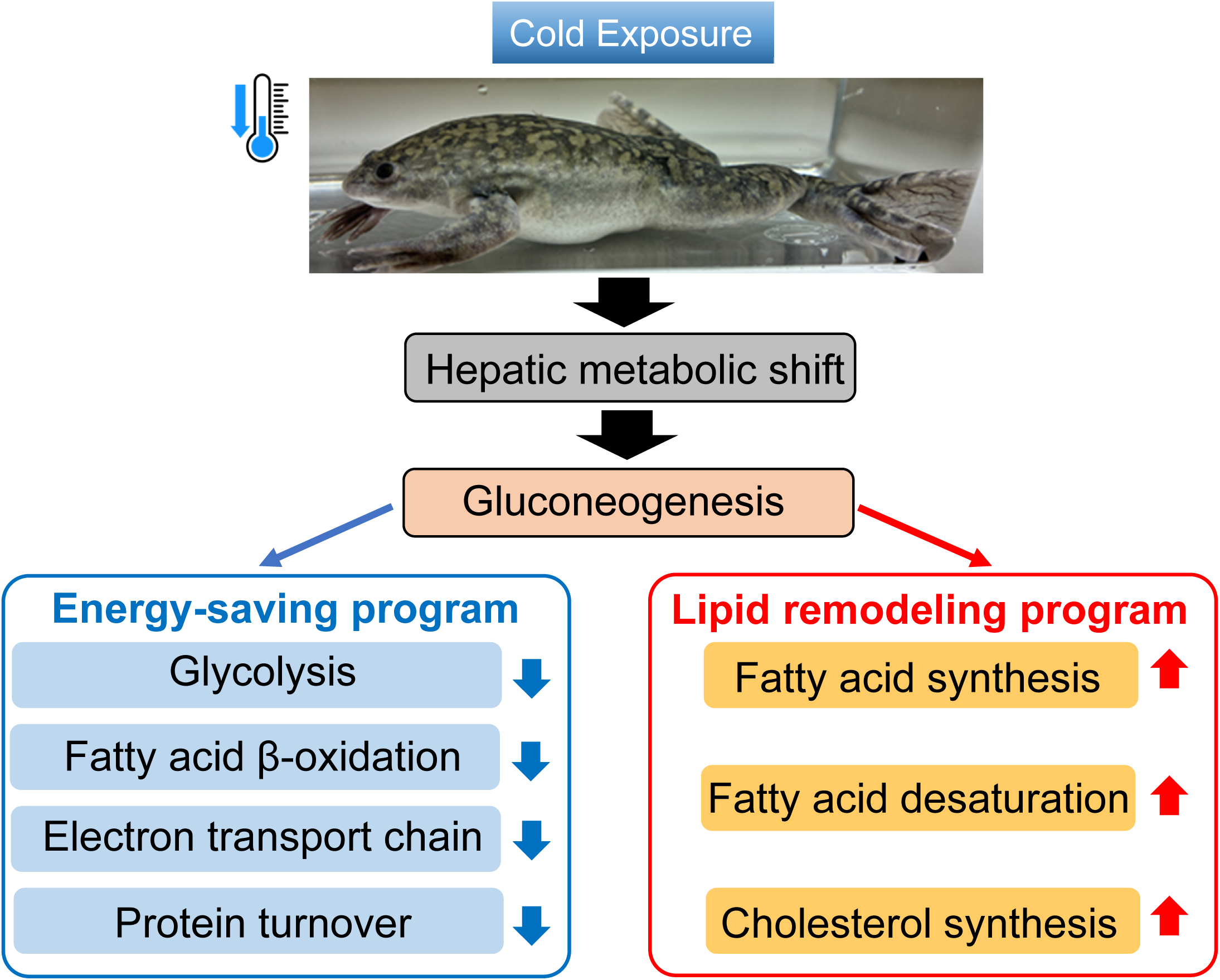
Graphical summary of cold-induced transcriptional reprogramming of hepatic metabolism in *X. laevis*.

## List of abbreviations

ANOVA: analysis of variance
ATP: adenosine triphosphate
cDNA: complementary DNA
DEGs: differentially expressed genes
DNase: deoxyribonuclease
ETC: electron transport chain
log₂FC: log₂ fold change
GO: Gene Ontology
KEGG: Kyoto Encyclopedia of Genes and Genomes
NEFA: non-esterified fatty acid
padj: adjusted *P* value
PBS: phosphate-buffered saline
qPCR: quantitative polymerase chain reaction
RIN: RNA integrity number
RNA-seq: RNA sequencing
SEM: standard error of the mean
TBARS: thiobarbituric acid reactive substances
TCA cycle: tricarboxylic acid cycle

## Declarations

### Ethics approval and consent to participate

All animal experiments were performed according to the Guide for the Care and Use of Laboratory Animals prepared by Hiroshima University (Higashi-Hiroshima, Japan).

### Consent for publication

Not applicable.

## Availability of data and materials

All raw sequencing data generated in this study have been deposited in the DDBJ Sequence Read Archive (DRA) under BioProject number PRJDB42163 (PSUB048229).

## Author’s contributions

Conceptualization: E.I.-U.; Methodology: E.I.-U., M.F.; Validation: E.I.-U., M.F.; Investigation: E.I.-U., M.S., M.F., N.S., Y.N., K.U.; Resources: M.S., H.O.; Writing - original draft: E.I.-U.; Writing - review & editing: E.I.-U., M.S., Y.N., K.U., H.O.; Visualization: E.I.-U.; Supervision: E.I.-U., H.O.; Project administration: E.I.-U.; Funding acquisition: E.I.-U.

## Acknowledgements

We thank the National BioResource Project (NBRP, Clawed Frogs/Newts) of MEXT.

---

**Table S1.**
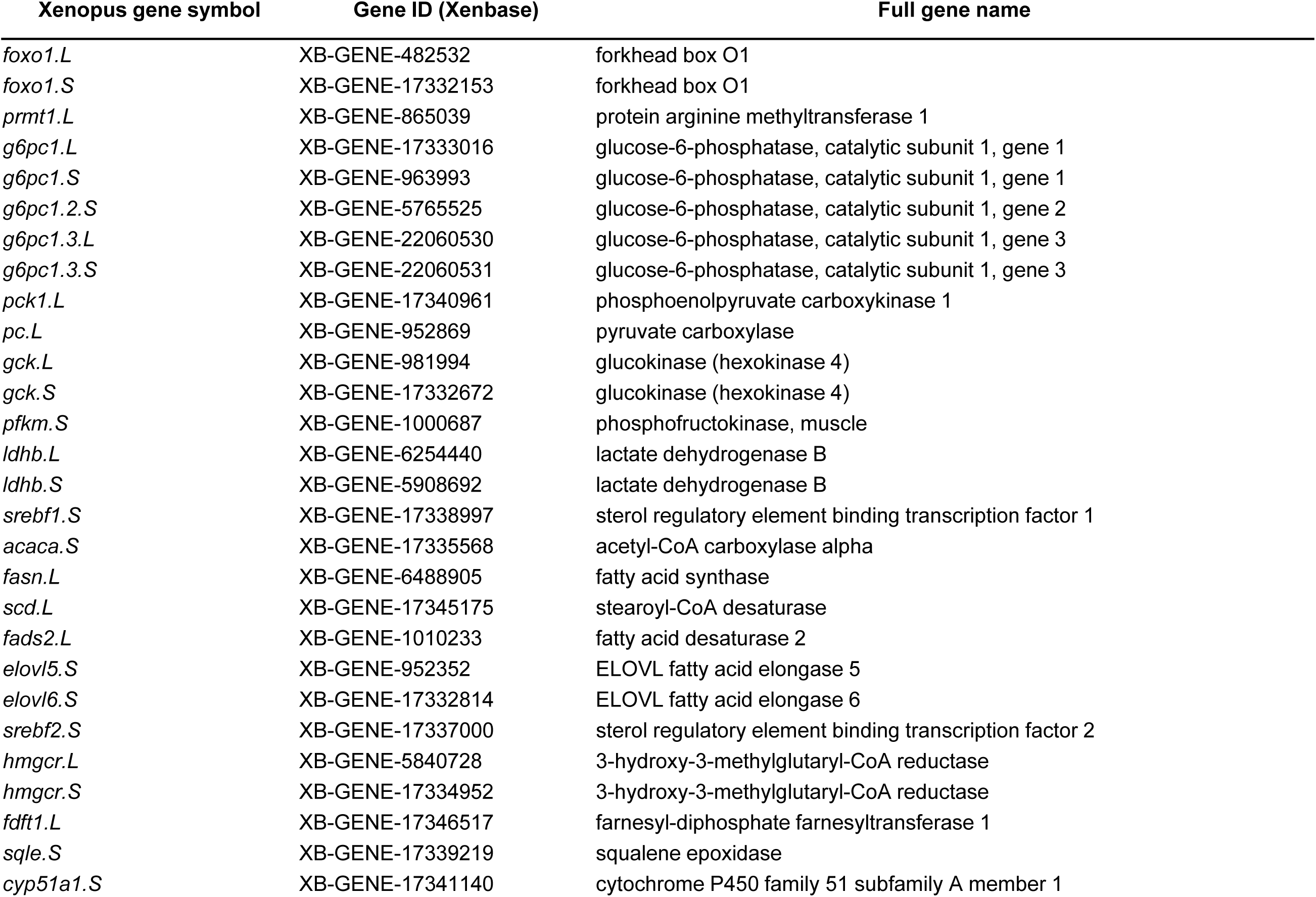
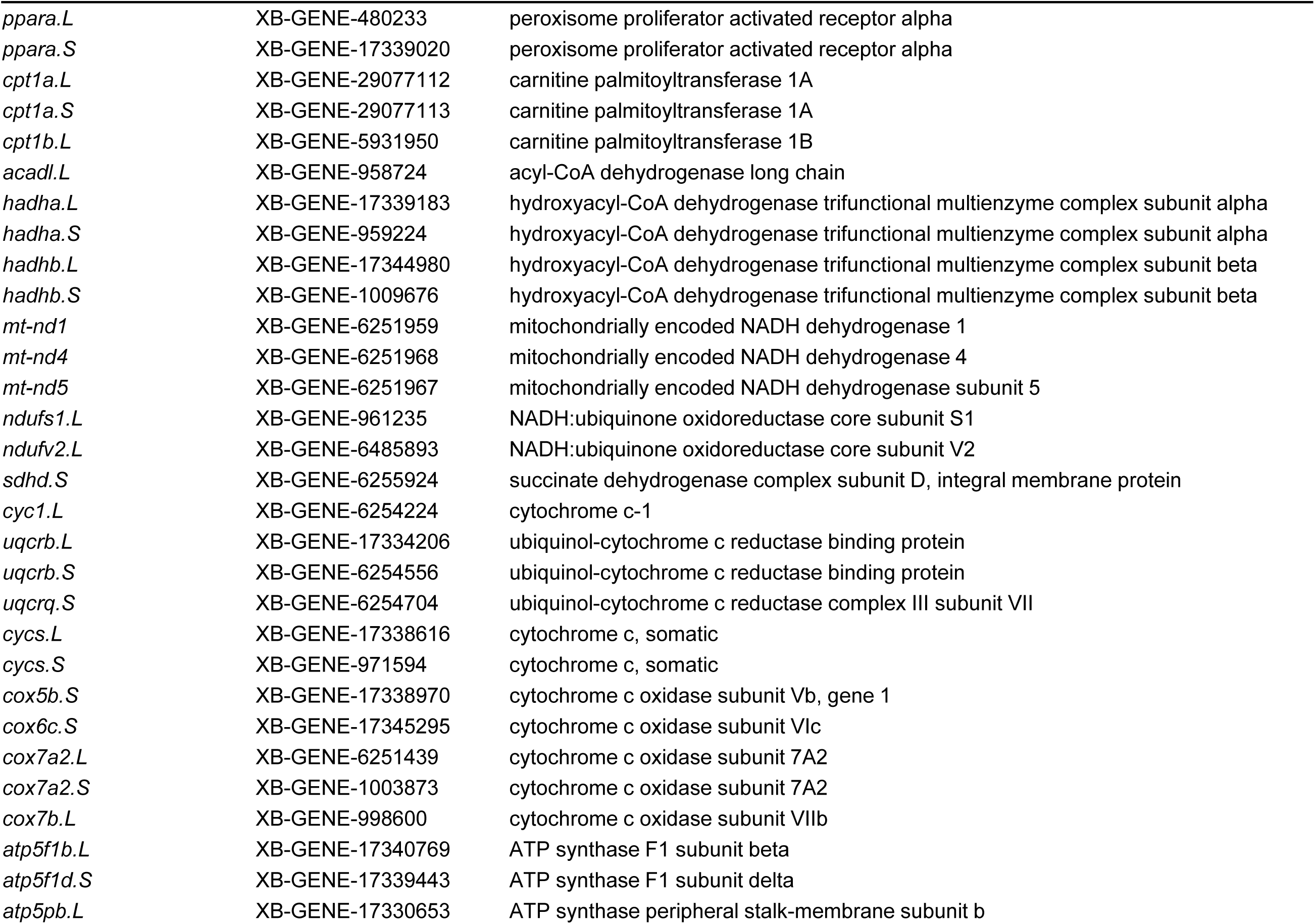
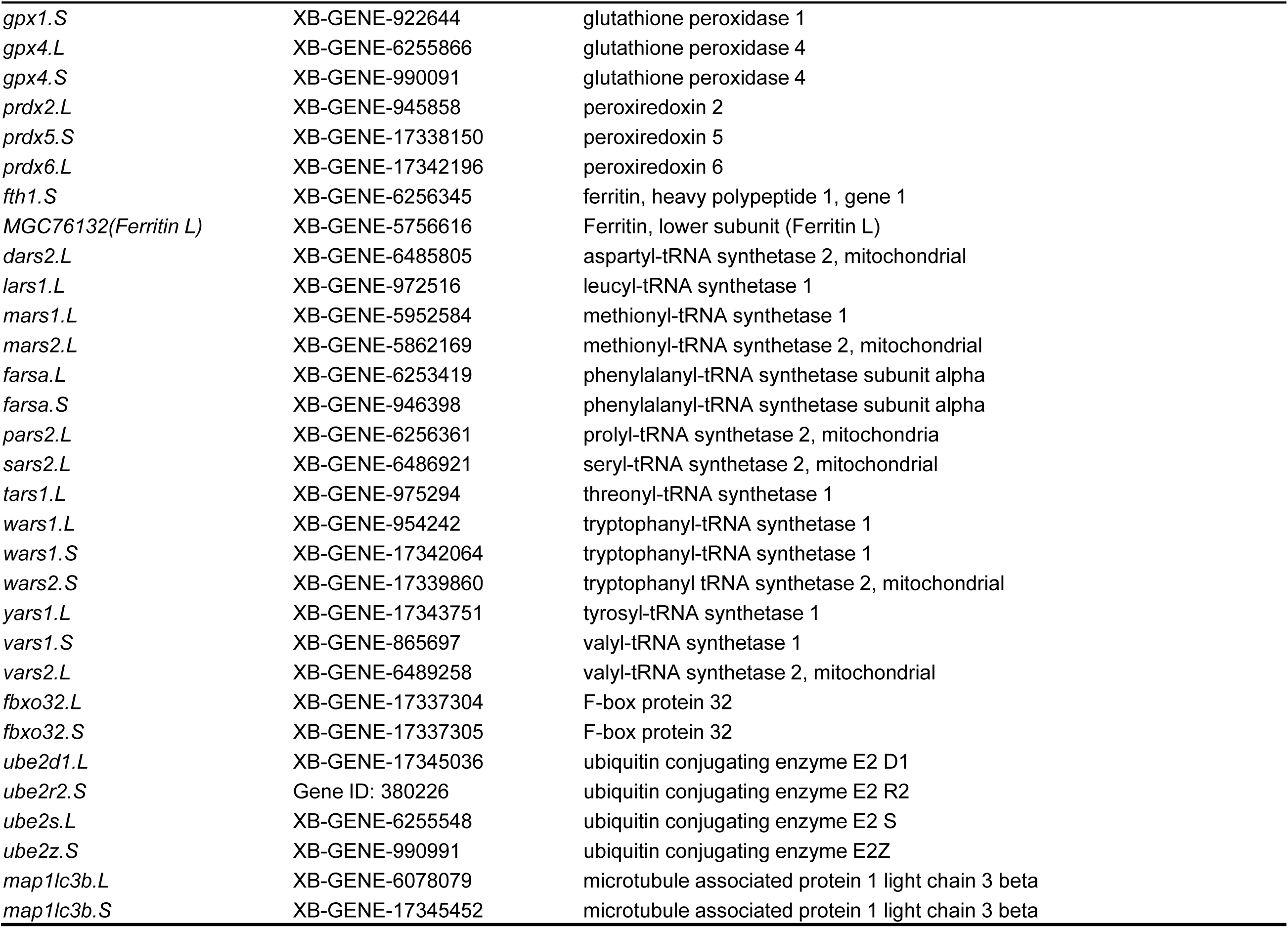
Gene abbreviation.

**Table S2.**
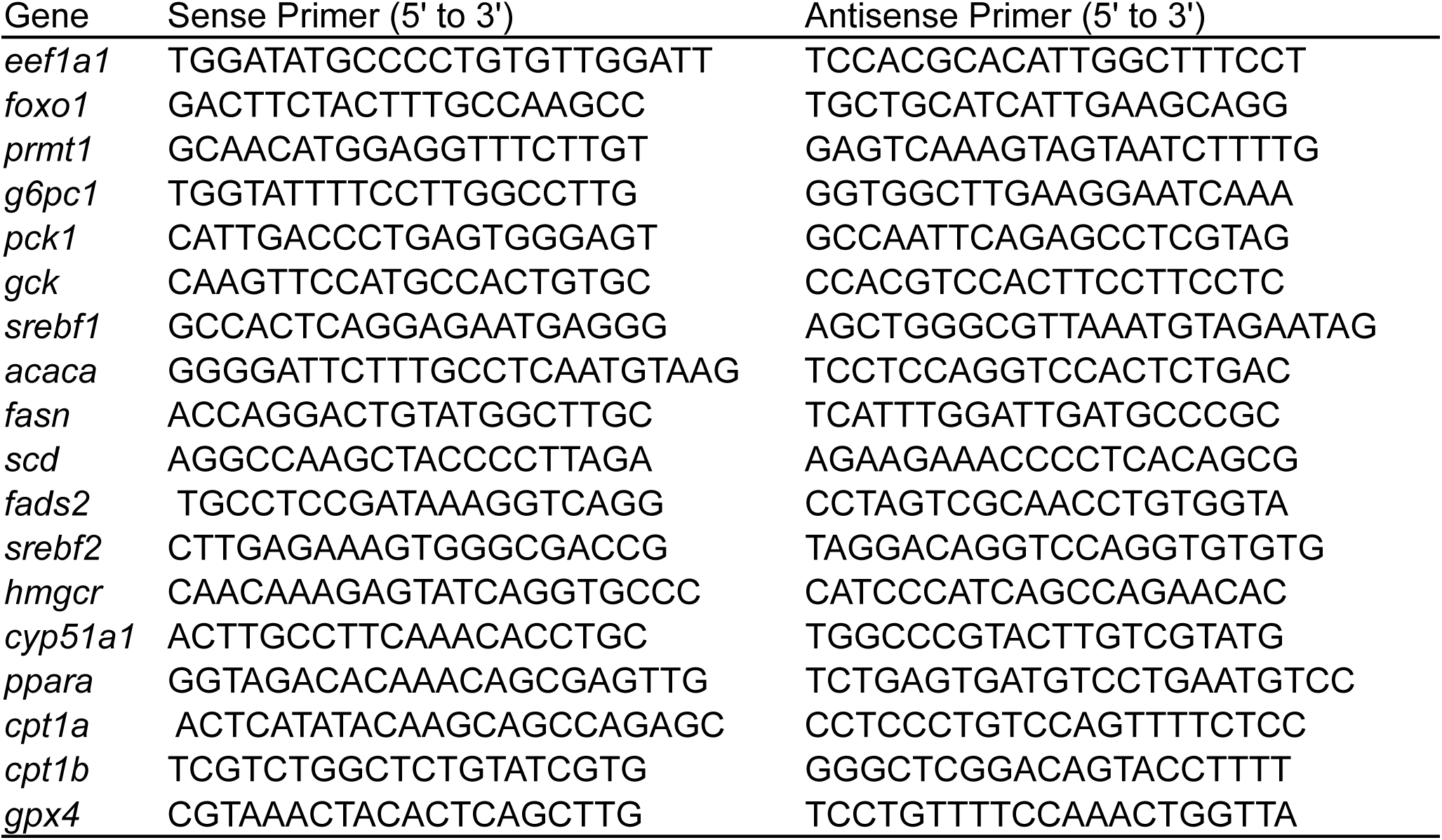
Sequences of oligonucleotide primers for qPCR.

## References

1. Storey KB, Storey JM. Biochemical adaption for freezing tolerance in the wood frog,Rana sylvatica. J Comp Physiol B. 1984;155:29–36. 10.1007/BF00688788.

2. Storey KB, Storey JM. Freeze tolerance and intolerance as strategies of winter survival in terrestrially-hibernating amphibians. Comp Biochem Physiol A Physiol. 1986;83:613–7. 10.1016/0300-9629(86)90699-7.

3. Storey KB, Storey JM. Molecular Physiology of Freeze Tolerance in Vertebrates. Physiol Rev. 2017;97:623–65. 10.1152/physrev.00016.2016.

4. Al-attar R, Wijenayake S, Storey KB. Metabolic reorganization in winter: Regulation of pyruvate dehydrogenase (PDH) during long-term freezing and anoxia. Cryobiology. 2019;86:10–8. 10.1016/j.cryobiol.2019.01.006.

5. Costanzo JP, Reynolds AM, Do Amaral MCF, Rosendale AJ, Lee RE. Cryoprotectants and Extreme Freeze Tolerance in a Subarctic Population of the Wood Frog. PLoS ONE. 2015;10:e0117234. 10.1371/journal.pone.0117234.

6. Alshwairikh YA, Skelly DK. The transcriptomics and differential gene expression of freezing and thawing in a freeze-tolerant vertebrate. BMC Genomics. 2026;27:167. 10.1186/s12864-026-12535-y.

7. Nagasawa K, Tanizaki Y, Okui T, Watarai A, Ueda S, Kato T. Significant modulation of the hepatic proteome induced by exposure to low temperature in *Xenopus laevis*. Biol Open. 2013;2:1057–69. 10.1242/bio.20136106.

8. Do Amaral MCF, Frisbie J, Crum RJ, Goldstein DL, Krane CM. Hepatic transcriptome of the freeze-tolerant Cope’s gray treefrog, Dryophytes chrysoscelis: responses to cold acclimation and freezing. BMC Genomics. 2020;21:226. 10.1186/s12864-020-6602-4.

9. De Mendoza D, Pilon M. Control of membrane lipid homeostasis by lipid-bilayer associated sensors: A mechanism conserved from bacteria to humans. Prog Lipid Res. 2019;76:100996. 10.1016/j.plipres.2019.100996.

10. Wu G, Baumeister R, Heimbucher T. Molecular Mechanisms of Lipid-Based Metabolic Adaptation Strategies in Response to Cold. Cells. 2023;12:1353. 10.3390/cells12101353.

11. Storey KB, Storey JM. Metabolic rate depression in animals: transcriptional and translational controls. Biol Rev Camb Philos Soc. 2004;79:207–33. 10.1017/S1464793103006195.

12. Van Breukelen F, Martin SL. Translational initiation is uncoupled from elongation at 18℃during mammalian hibernation. Am J Physiol Regul Integr Comp Physio. 2001;281:R1374–9. 10.1152/ajpregu.2001.281.5.R1374.

13. Storey KB, Storey JM. Metabolic rate depression: the biochemistry of mammalian hibernation. Adv Clin Chem. 2010;52:77–108.

14. Van Breukelen F, Martin SL. The Hibernation Continuum: Physiological and Molecular Aspects of Metabolic Plasticity in Mammals. Physiology. 2015;30:273–81. 10.1152/physiol.00010.2015.

15. Carey HV, Andrews MT, Martin SL. Mammalian Hibernation: Cellular and Molecular Responses to Depressed Metabolism and Low Temperature. Physiol Rev. 2003;83:1153–81. 10.1152/physrev.00008.2003.

16. Vucetic M, Stancic A, Otasevic V, Jankovic A, Korac A, Markelic M, et al. The impact of cold acclimation and hibernation on antioxidant defenses in the ground squirrel (Spermophilus citellus): An update. Free Radic Biol Med. 2013;65:916–24. 10.1016/j.freeradbiomed.2013.08.188.

17. Sone M, Mitsuhashi N, Sugiura Y, Matsuoka Y, Maeda R, Yamauchi A, et al. Identification of genes supporting cold resistance of mammalian cells: lessons from a hibernator. Cell Death Dis. 2024;15:685. 10.1038/s41419-024-07059-w.

18. Session AM, Uno Y, Kwon T, Chapman JA, Toyoda A, Takahashi S, et al. Genome evolution in the allotetraploid frog Xenopus laevis. Nature. 2016;538:336–43. 10.1038/nature19840.

19. Sugasawa T, Ono S, Yonamine M, Fujita S, Matsumoto Y, Aoki K, et al. One Week of CDAHFD Induces Steatohepatitis and Mitochondrial Dysfunction with Oxidative Stress in Liver. Int J Mol Sci. 2021;22:5851. 10.3390/ijms22115851.

20. Zhang T, Jia L, Niu Z, Li X, Men S, Jiang L, et al. Comparative transcriptomic analysis delineates adaptation strategies of Rana kukunoris toward cold stress on the Qinghai-Tibet Plateau. BMC Genomics. 2024;25:363. 10.1186/s12864-024-10248-8.

21. Matsumoto M, Pocai A, Rossetti L, DePinho RA, Accili D. Impaired Regulation of Hepatic Glucose Production in Mice Lacking the Forkhead Transcription Factor Foxo1 in Liver. Cell Metab. 2007;6:208–16. 10.1016/j.cmet.2007.08.006.

22. Rehman S, Hadj-Moussa H, Hawkins L, Storey KB. Role of FOXO transcription factors in the tolerance of whole-body freezing in the wood frog, Rana sylvatica. Cryobiology. 2023;110:44–8. 10.1016/j.cryobiol.2022.12.018.

23. Fedotcheva NI, Litvinova EG, Kamzolova SV, Morguno IG, Amerkhanov ZG. Mitochondrial metabolites in tissues as indicators of metabolic alterations during hibernation. Cryo Lett. 2010;31:392–400.

24. Brooks SPJ, Storey KB. Mechanisms of glycolytic control during hibernation in the ground squirrel Spermophilus lateralis. J Comp Physiol B. 1992;162. 10.1007/BF00257932.

25. Shekhovtsov SV, Bulakhova NA, Tsentalovich YP, Zelentsova EA, Meshcheryakova EN, Poluboyarova TV, et al. Metabolomic Analysis Reveals That the Moor Frog Rana arvalis Uses Both Glucose and Glycerol as Cryoprotectants. Animals. 2022;12:1286. 10.3390/ani12101286.

26. Hazel JR. Thermal Adaptation in Biological Membranes: Is Homeoviscous Adaptation the Explanation? Annu Rev Physiol. 1995;57:19–42. 10.1146/annurev.ph.57.030195.000315.

27. Nozawa Y. Adaptive regulation of membrane lipids and fluidity during thermal acclimation in Tetrahymena. Proc Jpn Acad Ser B Phys Biol Sci. 2011;87:450–62. 10.2183/pjab.87.450.

28. Staples JF. Metabolic suppression in mammalian hibernation: the role of mitochondria. J Exp Biol. 2014;217:2032–6. 10.1242/jeb.092973.

